# BBQ methods: Streamlined workflows for Bacterial Burden Quantification in infected cells by confocal microscopy

**DOI:** 10.1101/2023.10.01.560379

**Authors:** Jacques Augenstreich, Michael Shuster, Yongqiang Fan, Zhihui Lyu, Jiqiang Ling, Volker Briken

**Affiliations:** Department of Cell Biology and Molecular Genetics, University of Maryland, College Park, MD 20742 USA; College of Life and Health Sciences, Northeastern University, Shenyang 110819, People’s Republic of China

## Abstract

Accurate quantification of bacterial burden within macrophages, termed Bacterial Burden Quantification (BBQ), is crucial for understanding host-pathogen interactions. Various methods have been employed, each with strengths and weaknesses. This article addresses limitations in existing techniques and introduces two novel automated methods for BBQ within macrophages based on confocal microscopy data analysis. The first method refines total fluorescence quantification by incorporating filtering steps to exclude uninfected cells, while the second method calculates total bacterial volume per cell to mitigate potential biases in fluorescence-based readouts. These workflows utilize PyImageJ and Cellpose software, providing reliable, unbiased, and rapid quantification of bacterial load. The proposed workflows were validated using Salmonella enterica serovar Typhimurium and Mycobacterium tuberculosis models, demonstrating their effectiveness in accurately assessing bacterial burden. These automated workflows offer valuable tools for studying bacterial interactions within host cells and provide insights for various research applications.

## Introduction

Accurate Bacterial Burden Quantification (BBQ) within macrophages plays a crucial role in understanding host-pathogen interactions; for example, in evaluating the efficacy of antimicrobial responses, or in studying the role of host or bacterial factors in pathogenesis. Several approaches have been employed to assess intracellular bacterial load, each with its own strengths and weaknesses. Existing methods include colony forming unit (CFU) count from cell lysate or different microscopy-based approaches including manual counting of bacteria per cell, total fluorescence or integrated density of fluorescence per field of view, bacterial area per cell or luciferase luminescence quantification. While these techniques have provided valuable insights, they often present limitations in terms of precision, automation, and unbiased quantification.

CFU count has long been a standard method for estimating relative bacterial load in a population of infected cells (Sutton, 2012; Jiang *et al*., 2021; Boamah *et al*., 2023; Mittal *et al*., 2023). It involves plating dilutions of lysates from infected macrophages on agar plates, allowing bacterial colonies to grow, and subsequently counting the colonies. Although CFU count provides a quantitative measure of bacterial burden, it requires time-consuming culturing steps and may underestimate the true bacterial load due to variations in bacterial growth conditions and recovery rates. This is particularly true in the case of mycobacteria, which can take 1-3 weeks to display a reliable colony count due to their slow growth (Welch *et al*., 1993). Another complication is that mycobacteria tend to aggregate and therefore rarely appear as single bacilli in the lysates of infected cells and these aggregates will appear as a single colony thus leading to an underestimation of the bacterial burden. In addition, the CFU counting itself can introduce human error that increases with the amount of samples to analyze (Sutton, 2012).

The use of bioluminescent bacteria and the luminescence readout can provide a precise but relative quantification of viable bacteria during macrophages infection. The main limitations are the access to strains expressing the reporter gene and the fact that the light emission is not compatible with multiplex microscopy detection. In addition, the expression of luciferase by the transfected bacteria may vary and/or be lost over time.

In general, microscopy provides a direct observation of the bacteria in the sample, so it gives a more precise and reliable quantification. This approach has been widely used for mycobacterial infection of macrophages as it provides a more rapid evaluation of bacterial burden compared to CFU count (Lerner *et al*., 2017; Mahamed *et al*., 2017; Aylan *et al*., 2023; Golovkine *et al*., 2023; Malaga *et al*., 2023; Raykov *et al*., 2023). The main limitation is to have a bacterial strain constitutively expressing a fluorescent marker, but either mycobacterial expression plasmids for fluorescence proteins or already transfected strains are now easily available. Several methods are described in the literature to quantify bacterial burden by microscopy. First, manual counting of bacteria per cell offers a direct assessment of bacterial burden at the single-cell level. This method involves visually inspecting microscopy images and manually enumerating the bacteria residing within each macrophage (Payros *et al*., 2021). While it allows for relatively precise quantification, manual counting is labor-intensive, subject to human error and bias, and impractical for analyzing large datasets. Additionally, it can be challenging to distinguish between closely clustered bacteria frequently encountered when working with mycobacteria for example. The quantification of the bacterial area per cell is another approach recently employed to estimate bacterial burden (Aylan *et al*., 2023). By segmenting bacteria within macrophages and measuring the area occupied by bacterial fluorescence, it is possible to assess the relative bacterial load per cell. While this method offers good insight into the intracellular bacterial distribution, it does not account for variations in bacterial size or the three-dimensional architecture of infected cells. Quantifying bacterial burden based on total fluorescence or integrated density of fluorescence per field of view offers a precise and relative quantitation approach (Golovkine *et al*., 2023; Malaga *et al*., 2023). By summing the fluorescence intensity of all pixels within a defined region, this method provides a measure of the relative bacterial load compared to the background fluorescence of an uninfected cell. It is thus very suitable to monitor the growth or survival of bacteria between different infection conditions for example. However, the non-specific background fluorescence or auto-fluorescence from the host cell can introduce variability and affect the accuracy of quantification.

To address these limitations, we propose two novel automated methods for BBQ within macrophages based on image analysis of confocal microscopy data. We first propose an improvement of the existing methods to collect the total fluorescence in infected cells (Lerner *et al*., 2017; Payros *et al*., 2021), by adding some filtering steps to exclude uninfected cells from the quantification, and by automatizing the process into the PyImageJ environment. Alternatively, we introduce a new method of quantification by calculating the total bacterial volume per cell, to circumvent potential biases in fluorescence-based readouts.

## Workflows – introduction

The goal of these workflows is to provide a reliable, unbiased and rapid method to quantify the bacterial load in infected cells. The example given here are RAW264.7 macrophages infected with YFP expressing strain of *Salmonella enterica* serovar Typhimurium (S.Tm). Before acquisition, the infected cells were fixed at the desired time post-infection and stained with DAPI (see material and methods section). Any other type of nuclei stain is expected to perform as well as DAPI.

The base image material used for this workflow are multi-channel, z-stack images acquired by confocal microscopy. The optimal organization of data; images and folders, for both workflows is illustrated in figure 1A. Each individual experiment is in their own folder or “parent folder”. The eventual multiple conditions would be in subfolders, containing the different fields of views or technical replicates. The workflows are presented as code contained in jupyter notebooks and can be adapted to many different types of images and indexing.

**Figure 1:**
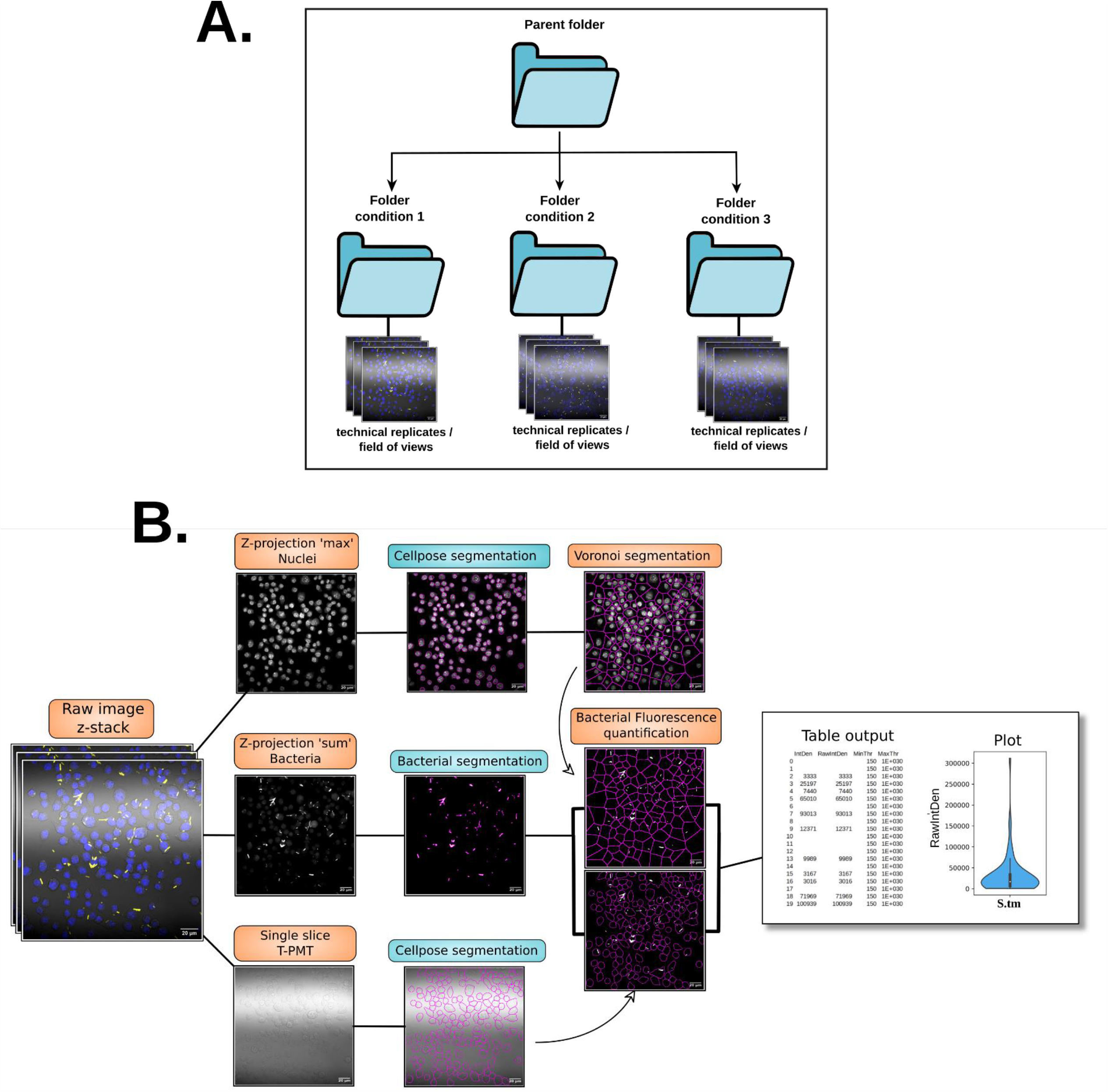
Diagram of the workflow for BBQ by total fluorescence quantification. (A) Representation of the optimal data organization for the workflows. (B) Diagram of the workflow with representative microscopy images from each steps. The “table output” and “plot” are representative of the quantification process.

For the image analysis, the automated workflows were implemented using the recently released ‘PyImageJ’ library, that allows the use of ImageJ/Fiji into a Python environment (Rueden *et al*., 2022). Detailed documentation for the installation and its use are described here (https://py.imagej.net/en/latest/). This library allows the combined use of Fiji and python that permit the image treatment on ImageJ and simultaneous use of python-based segmentation or tracking software under the same ecosystem.

For nuclei and cell segmentation, we used the deep learning-based segmentation software Cellpose (Stringer and Pachitariu, 2022). This software has the major advantage of facilitating training of custom segmentation models, thus that can be adapted to any cell type where a difference in cell morphology can decrease segmentation efficiency. Cellpose can also be invoked using python commands, so side by side with Fiji in the same environment. We took advantage of this flexibility and interoperability to propose nearly fully automated image analysis workflows that can be adapted to multiple types of images. We propose one workflow to quantify the total bacterial fluorescence intensity per cell and a second workflow for the total bacterial volume per cell.

The code and process are described in detail in the associate notebooks available on GitHub (https://github.com/jaugenst/BBQ/). Each step of the workflow is represented in the notebooks as a cell or block of code that can be ran sequentially or all at once if desired and the proper data pre-processing performed.

## 1. Workflow – Quantification of total bacterial fluorescence

All the steps described below are illustrated in the diagram in figure 1B. The code of the workflow is contained in the notebook “bacterial_burden”.

### Step 1: Nuclei channel isolation and Cellpose segmentation

To isolate signals from single cells, the Voronoi network analysis method is used and relies first on the detection of nuclei from the image. To this end, the nuclei channel is selected from the original z-stack image and is compressed into a smoothed z-projection ‘max’, and finally saved. The nuclei signals are then detected and outlined using Cellpose. This software can provide as an output the coordinates for Regions Of Interests (ROI) in a text file that can be converted in ROIs directly in ImageJ/Fiji.

### Alternative Step 1

If the use of DIC or transmitted light channel is possible, these channels can be used for cell segmentation which would outline the cells more precisely compared to Voronoi segmentation. Cellpose can be used to generate a model able to segmentate the cells from this type of channel (figure 1B) and as previously showed, can produce ROIs as an output directly usable in ImageJ/Fiji (Augenstreich *et al*., 2022). This overrides the need for a subsequent Voronoi segmentation and from this step it is possible to go directly to step 3.

### Step 2: Voronoi segmentation

The ROIs of the segmented nuclei in step 1 are used to perform the Voronoi segmentation in Fiji and adapted from a method described elsewhere (Payros *et al*., 2021) and that was also adapted from another study (Lerner *et al*., 2017). Briefly, it involves particle detection to collect ROIs’ centroids, and then perform the Voronoi segmentation. As an improvement, and to ensure proper and automatic ROI centroid detection, the ROIs are first reduced in size by 1 pixel to make sure that the subsequent particle detection can detect single objects. Indeed, 2 objects separated by only one pixel will frequently be detected as one, and watershed function is not always successful at separating objects. The ROIs are also used to remove the background signal and are filled to facilitate the transformation into a binary image. After the latter is performed, the centroids of nuclei are collected and used the Voronoi segmentation is performed, and ROIs are created from each zone given by the segmentation.

### Step 3: Z-projection, and bacterial detection

To measure the total bacterial fluorescence per macrophage, the intracellular bacteria must be detected and their signal isolated in order to be measured. To do so, the bacterial channel is isolated, and a z-projection ‘sum’ is created and saved. This image is duplicated, smoothed, thresholded and transformed into a binary image. The bacteria are then detected, and ROIs are created around them and saved. This new ROI set is then applied back to the original z-projection image to clear any signal outside the bacteria.

### Step 4: total bacterial fluorescence quantification and infectivity measurement

The ROIs from the Voronoi segmentation are called and applied on the cleared bacteria z-projection and the total fluorescence (RawIntDen) is collected in each cell. The results table obtained is automatically exported as a CSV file and saved and will display all the cells as a list of numbers and their corresponding total bacterial fluorescence value which represent the bacterial burden (figure 1B). In the absence of bacteria, the cell will have an N/A value in the Fiji results table and no value displayed in the CSV table (figure 1B). Thus, the ratio of the number of cells containing bacterial fluorescence compared to the total number of cells detected would give the percentage of infected cells / infectivity and can be easily calculated from the CSV file.

At the end of the workflow, the folders containing the different images of eventual different conditions of some experiment should now be populated with the images of isolated nuclei channel, the isolated bacterial channel, the cellpose output (png and txt), and a zip file containing the ROIs from the Voronoi segmentation, thus allowing to control the results of the different step if needed. Each folder should also contain a CSV file named after the folder name containing all the results of measurements from each image of this folder concatenated in this single file.

## 2. Workflow – Quantification of total bacterial volume per cell

We established an alternate method for the quantification of bacterial burden which we propose to be an improvement of measurement of bacterial burden based on total fluorescence intensity. We propose that relying on bacterial fluorescence alone for bacterial burden quantification is potentially biased by different parameters. The main bias is that this quantification is based upon on the assumption that the expression of fluorescent proteins remains constant over time. Nevertheless, we provide evidence that expression of fluorescence proteins driven by a constitutive promoter change when bacteria are in different environments (figure 3).

To circumvent this bias, we propose a workflow that aims to measure an estimation of the total bacterial volume inside the infected cells which is independent of fluorescence intensities. The base image material for this workflow is similar to the one described for the first workflow, as it is designed for a multi-channel, z-stack image of cells infected with fluorescent bacteria or your favorite microorganism. The main difference is about the data collection after either the Voronoi ROIs or the cell outlines from Cellpose are generated. Instead of collecting the raw integrated density of fluorescence on a z-projection sum of the bacterial channel, it’s the bacterial area that is measured directly on the z-stack and used for further calculation. From the area measurement on each slice of the z-stack, it is possible to calculate estimated partial bacterial volumes, by multiplying the area with the interval between two slices (figure 2A). Ultimately, the sum of the partial volumes will estimate the total bacterial volume per cell in µm^3^. A diagram illustrating the workflow is given in figure 2B. The code of the workflow is contained in the notebook “bacterial_burden_volume” in the GitHub repository.

**Figure 2:**
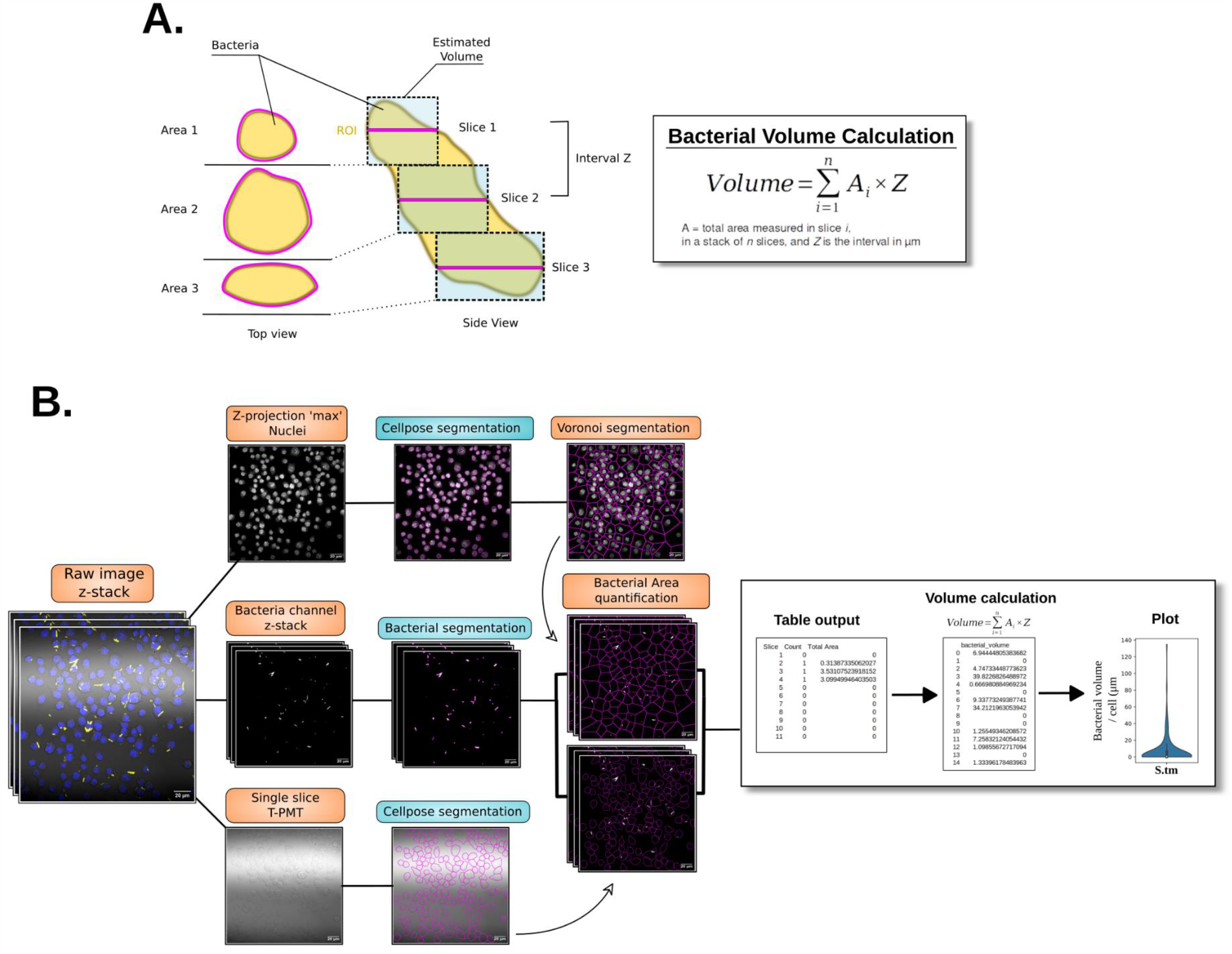
Diagram of the workflow for BBQ by total bacterial volume calculation. (A)Principle of bacterial volume calculation. The bacteria when imaged are horizontally sliced at a define interval Z. The area A for each slice is collected and the total volume calculation is performed following the equation in the right panel. (B) Diagram of the workflow with representative microscopy images from each steps. The “table output”, the “Volume calculation” table and the “plot” are also representative of the quantification process.

### Step 1: Nuclei channel isolation and Cellpose segmention

As decribed earlier, from the original z-stack image, the nuclei channel is selected and collapsed into a smoothed z-projection ‘max’, and finally saved. The nuclei signals are then detected and outlined using the Cellpose segmentation software. This software also allows to obtain Regions of Interests (ROI) from a text file that can be converted in ROIs directly in ImageJ/Fiji.

### Alternative Step 1

As explain above, if the use of DIC or transmitted light channel is possible, Cellpose is then used to segmentate the cells and produce ROIs as an output directly usable in ImageJ/Fiji. This overrides the need for a subsequent Voronoi segmentation and it is possible to go directly to step 3.

### Step 2: Voronoi segmentation

The detected nuclei in step 1 are used to perform the Voronoi segmentation in Fiji. The nuclei ROIs are first reduced in size by 1 pixel to make sure that for further particle detection can detect single objects. The ROIs are also used to remove the background signal and are filled to facilitate the transformation into a binary image. After the latter is performed, the nuclei centroids are collected and used the Voronoi segmentation is performed, and ROIs are created from each zone given by the segmentation.

### Step 3: Bacterial detection and measurement of bacterial area per slice

The bacterial channel is first isolated, smoothed, thresholded and transformed in a binary image. The bacteria are then detected by particle detection function, and ROIs are created around them on each slice and saved. While the ROIs are being created, the “analyze particle” function also allows for the simultaneous collection of data by choosing the “summarize” option. It displays a table showing how many particles were detected per slice and the associated total area. The collection of data is iterated on each Voronoi ROI and the final summary table is saved.

### Step 4: Bacterial Volume Calculation

The calculation of the volume for some regular shaped objects is the product of the area by the height. If the imaged object is irregular but sliced, it is possible to calculate partial volumes based on the slicing thickness of the object (here the z-stack step) and the different values of bacterial area along the z-stack (figure 2A). For our purpose of quantifying volume of bacteria an estimation of the total volume would be the sum of all the partial volume, that can be resumed in this equation:

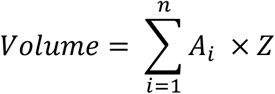

where A is the total area measured in the slice *i* in a stack of n slices, and Z is the interval in µm between 2 slices. The workflow will extract the summary table generated in step 3 and calculate the volume by first summing the total area values in 11 rows at a time (or number of slices in the z-stack). This sum of area is then multiplied by the distance between two slices. The product will be the bacterial volume in µm^3^. The calculation is automatically reiterated at every repetition of 11 rows but can be adapted for your own set of images and their corresponding slicing. The result of this calculation is stored in a new table and the calculation is reiterated for each summary table of each image and concatenated in the new table. Finally, this results table is automatically saved in the matching folder and named after the folder.

In the end of the workflow, the folders containing the different images of eventual different conditions of some experiment should now be populated with the isolated nuclei channel, the isolated bacterial channel, the Cellpose output (png and txt), and a zip file containing the ROIs from the Voronoi segmentation, thus allowing to control the results of the different step if needed. Each folder should also contain a series of CSV files containing the summary table of each image, and a CSV file named after the folder name containing all the volume calculation results from each image of this folder.

## Validation results and examples

We proposed the need for development of a method to quantify bacterial burden by determination of the total bacterial volume per cell was because of the potentiality that bacterial expression level of the fluorescent reporter gene could change depending on stresses encountered by bacteria even if the gene is regulated by a constitutive promotor. We tested this hypothesis using a *Salmonella enterica* serovar

Typhimurium (S.Tm) strain that constitutively expresses mCherry, a fluorophore typically known for its resistance to acidic pH (Doherty, Bailey and Lewis, 2010). We exposed S.Tm-mCherry to three different stresses that the bacteria are likely to encounter in a phagosome such as low pH, reactive oxygen species (ROS) or reactive nitrogen species (RNS). Single bacterial fluorescence analysis was performed as previously described (Lyu *et al*., 2023). We show that in an acidic pH, bacterial mCherry signal was lower than in a neutral pH (figure 3A). In contrast, mCherry expression was higher for bacteria cultivated in medium supplemented in H_2_O_2_ as an oxidative stress (figure 3A). In a similar fashion, a nitrosative stress induced by Spermine-NONOate (SPER/NO) increased fluorescence of mCherry compared to the control condition (figure 3A). The bacterial area was also measured and showed similar between the conditions which demonstrates that the fluorescence variation is not the result of a difference in bacteria (figure 3B). Overall, these results demonstrate that the environment could directly bias constitutive expression level of reporter genes. Consequently, this will affect the reliability of bacterial burden measurements using fluorescence readouts and possibly luminescence readouts used to quantify live bacteria (Andreu *et al*., 2010).

**Figure 3:**
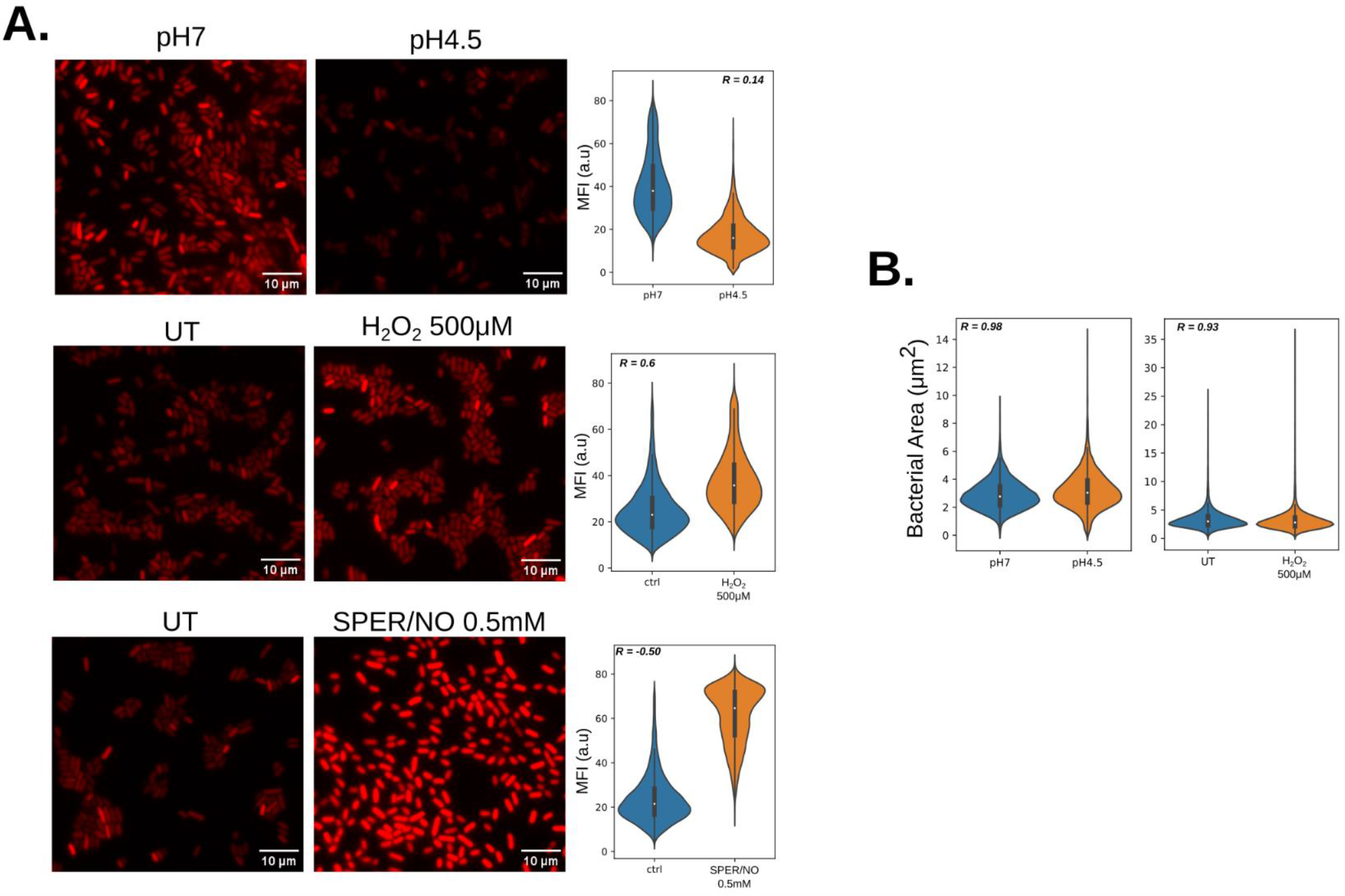
Differential expression of mCherry reporter gene in acidic or oxidative stress. (A)left panel; Representative image of bacteria incubated in normal LB (left column) or in LB at pH 4.5, LB+ H_2_O_2_ 100µM, or LB + SPER/NO 0.5mM for 16h. The bacteria were then imaged by wide-field fluorescence microscopy. Right panel; the bacteria were segmented using cellpose and the mean fluorescence intensity (MFI) was measured. The graph is showing the MFI values distribution from single bacterium from 3 biological replicates (*N*_pH7_ = 14607, *N*_pH5_ = 7414, *N*_UT_H2O2_= 12503, *N*_H2O2_ = 20100, *N*_UT_SPER/NO_ = 16103, *N*_SPER/NO_ = 18336. (B) Area quantification of single bacteria incubated indicated pH, left untreated (UT) or treated with 500µM H_2_O_2_ . The spearman R coefficient displayed was calculated between the frequency distribution of the 2 groups on each graphs.

Next, we assessed the precision of the bacterial volume calculation compared to the more classically used total fluorescence-based quantification. The two workflows were used side by side on images of RAW264.7 cells infected with S.Tm expressing YFP. To induce variation in the bacterial burden, the cells were pre-treated with cytochalasin-D that prevents phagocytosis, and thus, decreases bacterial burden. The direct cell segmentation by Cellpose from the Transmitted-light channel (T-PMT) was also chosen to exclude the extracellular bacteria that could still be visibly adherent to the cells (figure 4A).

**Figure 4:**
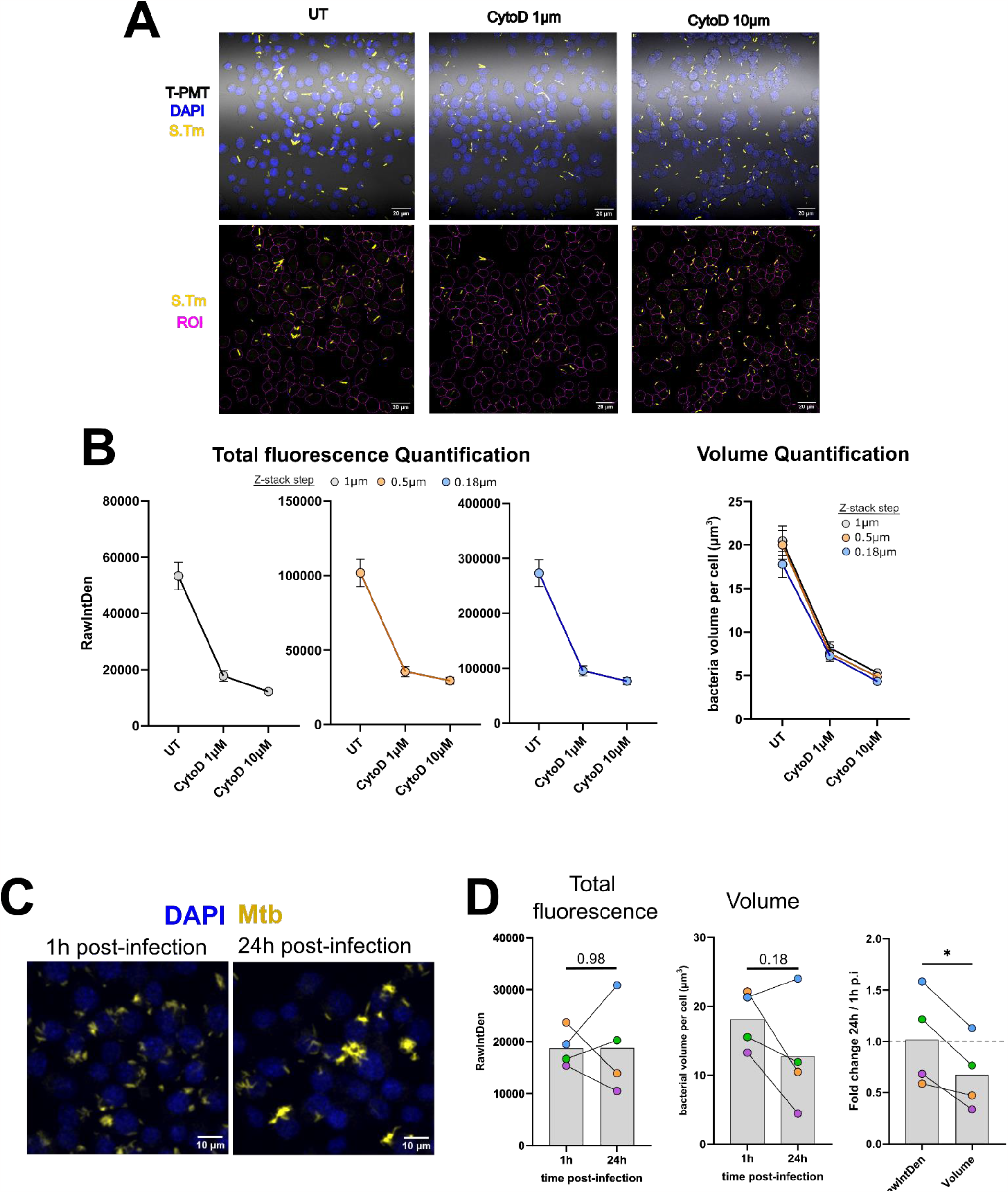
Total bacterial volume calculation performs similarly to total fluorescence measurement and can compensate from fluctuation in reporter expression. (A-B) RAW264.7 cells were treated with 1µM or 10µM cytochalasinD (CytoD) or left untreated for 1h, and were infected with YFP expressing S.Tm at MOI 10 for 30 min. The cells were then fixed, stained and mounted with DAPI and imaged by confocal microscopy. The same fields were images three times with a 10µm range, but with 11 slices (1µm interval), 21 clices (0.5µm interval) and 56 slices (0.18µm interval) (A) Representative images of RAW264.7 cells infected with S.Tm. The lower panel shows the bacterial fluorescence channel with the ROIs obtained from cellpose segmentation overlaid to illustrates the areas of the images where the fluorescence or the bacterial area were collected. (B) Quantification of total bacterial fluorescence or bacterial volume per cell, from images taken at 1µm, 0.5µm or 0.18µm distance interval between the slices of 10µm z-stacks. The results are representative from 3 fields of views per conditions. (C-D) RAW cells were infected with a DsRed-expressing strain of H37Rv (Mtb) for 1h at MOI 10, and incubated for 1h or 24h. At the designated timepoint, the cells were fixed, stained and mounted with DAPI and imaged by confocal microscopy. (C) Representative images of RAW cells infected with Mtb at 1h and 24h post infection. (D) total bacterial fluorescence quantification per cell (left) or total bacterial volume per cell (Middle). (Right) Comparison of fold change between 1h and 24h p.i. The points represent the mean value from 4 independent experiments. The statistical significance[para-merge-check

The cells were imaged at a fixed pinhole aperture of 1 airy unit, and the range of the z-stack was fixed at 10 µm. The same field were acquired with different steps between the slices, 1 µm, 0.5 µm that corresponds to the resolution in z at 1 airy unit, and 0.18 µm that corresponds to the distance that match Nyquist Sampling. As shown, the calculation of the bacterial volume proved to be similar, with about 7% of volume variation on average between the sampling methods (figure 4B). This is reasonable in the light of acquisition time where one must adjust between sampling precision and the number of fields of views acquired. Here even a sampling at 1 µm returned comparable accuracy in volume calculation compared to thinner sampling (figure 4B). This accuracy was conserved on the other condition tested where the cells were treated with cytochalasin D (CytoD) before getting infected. And indeed, the expected decrease of bacterial burden by CytoD treatment was observed at any sampling distance with a marginal difference. Finally, the volume calculation was compared to total fluorescence quantification (figure 4B). Both quantification methods detected the decrease in a similar fashion thus performed similarly to detect a decrease in bacterial burden. Given the similarity in trends of the different curves, these results also highlight that a slicing of 1 µm that could be considered as an under-sampling gave an excellent approximation of bacterial burden, for both methods of quantification.

In order to validate even further the bacterial volume workflow, we assessed the bacterial burden of RAW264.7 cells infected with *Mycobacterium tuberculosis*, a bacterium that can persist for long periods of time in macrophages (figure 4C-D). After infection, the progression of bacterial burden was analyzed 1 h and 24 h after infection. First, total fluorescence measurement showed great variability between independent experiments but on average didn’t detect any changes in bacterial burden (figure 4D). However, bacterial volume calculation showed a clear trend of a decreased bacterial burden at 24 h p.i. compared to 1 h p.i. on 3 out the 4 independent experiments (figure 4D). Indeed, observation of the micrographs indicate that the cells looked less loaded in bacteria but appeared brighter (figure 4C). The comparison of the fold changes at 24 h between the total fluorescence measurement and the volume measurement confirmed that the fluorescence readout systematically overestimated bacterial burden per cell. This data strongly suggests that the results obtained *in vitro* on S.Tm for variable expression of reporter genes due to environmental stresses (figure 3) are actually observed in macrophage infected with mycobacteria.

Overall, these results support the use of bacterial volume calculation for long term analysis of bacterial burden quantification rather than fluorescence-based read-outs. Other reporter gene models such as Luciferase genes expression, often rely on the same promotors and hence their expression levels are also likely to be affected by the bacterial environment. In our specific example of Mtb we did not test luminescence readouts, but commonly used luciferase reporter plasmids contain the same *hsp6*0-derived promoter that is present in our DsRed expression plasmid backbone pMan-1 (Manzanillo *et al*., 2012). Consequently, our results suggest that this luciferase reporter gene expression might also be affected by the intracellular environment, independently of the survivability or fitness of the bacteria.

In summary, we present automated workflows for the determination of bacterial burden in infected cells by total fluorescence quantification or bacterial volume calculation. This can allow accurate monitoring of bacterial growth in cells, or eventually bacterial clearance due to host cell response or drug treatment. It can also provide users with an easy measure of infectivity, that is a relevant readout for example for the study of mutant strains and their interaction with macrophages.

## Material and methods

### Software and installation

It is highly recommended to install any software and libraries required for this workflow in a virtual environment, using Anaconda, Pyvenv or environment modules. A detailed documentation of pyimagej is available online (https://py.imagej.net/en/latest/). To be noted that this workflow is not functional on MacOS as the “interactive mode” of initialization of Fiji is not compatible with the operating system. The workflow development was done in a jupyter notebook in the software jupyter-lab (https://jupyter.org/). Cellpose software was installed following the documentation and installation instructions available online (https://github.com/mouseland/cellpose). ChatGPT 3.5 (OpenAI) was occasionally used for code debugging and editing.

### Cell culture, bacterial culture

The cells used in this study were RAW-Lucia™-ISG (invivogen) or RAW264.7 cells. They were cultivated in DMEM medium (Gibco) supplemented with 10% heat-inactivated fetal bovine serum (Thermofisher). The day prior of the infection, the cells were detached using a cell scrapper and seeded into a 24 wells plate in fresh medium that was supplemented in M-CSF 40ng/mL for the case of infections with Mtb.

The first bacteria used for the study are *Salmonella enterica spp* Typhimurium (S.Tm) expressing YFP or mCherry. The S.Tm chromosomal integration of the yellow fluorescent protein (YFP) is constructed according to the method described here (Fan *et al*., 2019). Briefly, chloramphenicol-yellow fluorescent protein cassette (cat-P_tet_-yfp) along with sequences homologous to the target region (3095232 bp to 3095506 bp) was integrated into the S.Tm chromosome by induction of Red recombinase, and the positive clones were selected by chloramphenicol and verified by sequencing. The strains expressing mCherry were generated by transformation with pZS-P_*tet*_-mCherry-TGA-YFP-X-ECFP. The X stands for promoter regions of *cadBA, hmpA* and *katG* genes that are activated by low pH, nitrosative stress and oxidative stress respectively, and each construct was used in the stressful condition corresponding to their respective activation properties. The strains were cultivated in Luria-Bertani (LB) medium at 37°C. Oligonucleotides used for this study are reported in table S1.

The second bacterial strain used for this study is *Mycobacterium tuberculosis* H37Rv (ATCC) expressing DsRed, described here (Augenstreich *et al*., 2022). The bacteria were grown in liquid Middlebrook 7H9 medium supplemented with 10% oleic acid-albumin-dextrose catalase (OADC) growth supplement, 0.2% glycerol, 0.05% tween 80, and Zeocin 100µg/mL.

### Study of bacterial stress and mCherry fluorescence intensity

For this study, overnight cultures of S.Tm containing their respective pZS-Ptet-mCherry-TGA-YFP-X-ECFP were 1:100 diluted in either LB (pH 7.0), acidic LB (pH adjusted to 4.5 by HCl) or LB supplemented with H_2_O_2_ at 500µM (Fisher Scrientic), Spermine-NONOate at 0.5 mM (Thermo Scientific Chemicals) and grown for 24 h at 37 °C. Cultures were harvested by centrifugation and resuspended in 20 μl LB. 1 μl of the resulting cultures were placed on a 1.5% agarose LB pad on a 12-well slide. The bacteria were then imaged by wide-field fluorescence microscopy (see procedure below).

### Macrophages infection

For infections with S.Tm, the bacteria were pelleted and resuspended in PBS followed by measuring the OD_600_ of the suspension. Bacteria for the infection were opsonized by taking 45 µL of bacterial culture, mixing with 20 µL of normal mouse serum (Jackson ImmunoResearch) and 135 µL of DMEM with 10% normal FBS (Gibco) for 20 minutes. This was followed by adding an additional 600 µl of DMEM with 10% normal FBS, and then adding the desired bacteria to the cells at an MOI of 10. The plate was spun at 100 x g for 5 minutes then incubation at 37°C for 30 minutes. The cells were washed three times with PBS, fixed with PFA 4% for 1 hour at room temperature. The cells were washed 3 times in PBS and mounted with Prolong mounting medium with DAPI (Molecular probes).

For infections with Mtb, the bacteria were pelleted and washed in PBS-tween80 0.05%. The OD600 was measured, and the bacteria were added on the cells at an MOI of 10. The plate was then spun at 200 x g for 5 minutes and incubated for 1 hour. The cells were washed three times with PBS and incubated with fresh complete medium. At the designated time post-infection, the cells were then washed three times with PBS, fixed with PFA 4% for 1 hour at room temperature. Finally, the cells were washed 3 times in PBS and mounted with Prolong mounting medium with DAPI.

### Microscopy

For the bacterial imaging, the images were obtained using a BZ-X800 fluorescence microscope (Keyence) with a 100x oil objective. The single bacteria fluorescence was collected as performed before(Lyu *et al*., 2023). Briefly, the bacteria were segmented on the mCherry channel using Cellpose. The single bacteria mean fluorescence intensity and area in µm^2^ was then measured. The data collection was processed in high throughput using the pyimagej library in jupyter-lab.

Imaging on the fixed infected cells was performed using a Zeiss laser scanning confocal microscope LSM-800, equipped with two gallium arsenide phosphide photomultiplier tube (GaAsP-PMT) detectors and a transmitted light photomultiplier tube detector (T-PMT), using the 63x/NA1.4 Oil objective. Each field of view was acquired as a Z-stack ranging 10 µm with a 1 µm step unless specified otherwise.

### Data analysis, statistical analysis and data visualization

The graphics were generated using either the seaborn library on python, or GraphPad prism 10. The statistical comparisons were performed on GraphPad prism 10. For the comparative study of stress responses, the large datasets were analyzed by comparing the frequency distributions using Spearman’s Rank Correlation Coefficient calculation. For the time course study, the means from each experiment and the fold change were compared by paired t-test.

All the figures and drawings were made and assembled using Inkscape.

### Data availability

All the code used for the workflows is available on GitHub (https://github.com/jaugenst/BBQ/). A dataset is available for testing here (Augenstreich and Briken, 2023).

## Acknowledgments

We sincerely thank Dr Serge Mazeres and Marc Augenstreich for their constructive feedback on the method validation. This work was supported by the National Institute of Allergy and Infectious Diseases (Grant R01AI139492 to V.B., R35GM136213 to J.L.). V.B. and J.A. designed the experiments, and wrote the manuscript. V.B., J.A., M.S, Z.L., and J.L., edited and proofread the manuscript. J.A., Z.L. and M.S. performed experiments. J.A. analyzed the results and developed the workflows. Y.F. and Z.L. created the fluorescent S.Tm strains.

We declare no conflicts of interests

